# Predicting treatment outcome using kinome activity profiling in HER2+ breast cancer biopsies

**DOI:** 10.1101/2022.09.23.508980

**Authors:** Donna O. Debets, Erik L. de Graaf, Marte C. Liefaard, Gabe S. Sonke, Esther H. Lips, Anna Ressa, Maarten Altelaar

## Abstract

In this study, we measured the kinase activity profiles of 32 pre-treatment tumour biopsies of HER2-positive breast cancer patients. The aim of this study was to assess the prognostic potential of kinase activity levels to identify potential mechanisms of resistance and to predict treatment success of HER2-targeted therapy combined with chemotherapy. Indeed, our system-wide kinase activity analysis, based on targeted mass spectrometry measurement of kinase activation loops, allowed us to link kinase activity to treatment response. Overall, high kinase activity in the HER2-pathway was associated with good treatment outcome. Furthermore, we found eleven kinases differentially regulated between treatment outcome groups. Amongst those, well-known players in therapy resistance were found, such as p38a, ERK and FAK, as well as a potential new player in drug resistance, namely MARK. Lastly, we defined an optimal signature of four kinases in a multiple logistic regression diagnostic test for prediction of treatment outcome (AUC=0.926). This kinase signature showed high sensitivity and specificity, indicating its potential as predictive biomarker for treatment success of HER2-targeted therapy.

## Introduction

Invasive breast cancer (IBC) is a highly heterogeneous disease, which is classified in subtypes with distinct molecular signatures (Hu et al. 2006). The disease course, survival rate and treatment strategy is highly dependent on the subtype. Around 15% of all breast cancer cases overexpress the human epidermal growth factor receptor 2 (ERBB2 or HER2), and is therefore referred to as HER2+ (HER2 positive) breast cancer. This subtype is more aggressive and typically has a poor treatment outcome (Harbeck et al. 2019; Andrulis et al. 1998; Owens, Horten, and Da Silva 2004). The development of antibodies blocking the HER2 receptor (Trastuzumab, TTZ, and Pertuzumab, PTZ) have improved the clinical outcome of HER2+ patients profoundly (Carter et al. 1992; Michailidou, Trenz, and de Wilde 2019; 2019; Owens, Horten, and Da Silva 2004; Schneeweiss et al. 2013). Consequently, these drugs have found their way to standard of care (Swain et al. 2015; 2013).

Treatment resistance against HER2-inhibition (both primary and acquired) is observed frequently (Pernas and Tolaney 2019). The development of treatment resistance has been extensively studied in the last decade resulting in the discovery of a multitude of different resistance mechanisms (Pernas and Tolaney 2019; Rimawi, Schiff, and Osborne 2015). Despite these discoveries, predictive biomarkers for treatment success are still absent, hampering effective clinical decision-making.

Many of the proposed resistance mechanisms evolve around rewiring of cellular signalling pathways. This can be either due to activation of alternative survival routes bypassing the HER2-pathway or the re-activation of HER2 and/or downstream signalling nodes via compensatory, redundant or mutated signalling molecules. The myriad of potential escape mechanisms demonstrates the need for patient-specific treatment strategies and hence appropriate patient stratification. However, the lack of effective diagnostic tools to pinpoint the rewiring mechanisms in a case-by-case fashion in a clinically relevant setting hampers biomarker identification and development of precision medicine.

Several strategies exist to measure rewiring of cellular signalling pathways; however, none of them provide all the relevant information. Since cellular signalling is heavily dependent on rapid and reversible protein phosphorylation (P Cohen 2001; Philip Cohen 2002), gene-, transcript- and protein-based analysis are insufficient. These techniques provide insights into pathway alterations, yet fail to pinpoint pathway activation. Phosphoprotein analysis, such as phosphoprotein-specific western blots and discovery phosphoproteomics, provide a more detailed view on which kinase substrates are activated within a pathway. Antibody-based techniques are generally very sensitive, yet their use is restricted by the limited availability of phosphosite-specific antibodies. Furthermore, due to its poor multiplexing capabilities antibody-based studies are generally focussed on a few phosphosites, providing limited pathway coverage and only allow for hypothesis-driven research. Phosphoproteomics-based techniques provide a more system-wide view, covering multiple signalling pathways in a single analysis, making it ideal to monitor signal transduction rewiring and to find tumour resistance mechanisms. However, determination of the exact kinase responsible for pathway rewiring is hampered by information-bias of kinase-substrate pairs and kinase motifs.

To overcome these limitations, a novel type of phosphoproteomics technology was developed previously, which allows for the direct measurement of kinase and pathway activity in a high-throughput and precise manner, covering a large set of kinases (currently, one third of the total kinome) (Schmidlin et al. 2019). This technology, QuantaKinome™, is based on measurement of the phosphorylation of kinase activation loops (T-loops), which in the majority of cases is a direct proxy for kinase activity (Nolen, Taylor, and Ghosh 2004). The precise quantification of such a large panel of kinase activities provides a comprehensive understanding of pathway activation and cellular rewiring in a simple, fast and precise way. Furthermore, since this approach does not rely on prior knowledge on substrate phosphorylation as surrogate for kinase activity, it allows for the discovery of the role of ‘dark’ or understudied kinases. Lastly, the unbiased system-wide approach grants the possibility of hypothesis-generating discovery studies.

In the current study, we applied this kinase activation loop assay to 32 treatment-naive HER2+ breast cancer biopsies to identify kinases and pathways linked to treatment success and to improve patient stratification. We found that increased HER2-pathway signalling was associated with treatment success. Furthermore, we identified 11 kinase activation states that were differentially regulated between treatment outcome groups. Lastly, we defined a panel of 4 kinases that were predictive of treatment outcome; the high specificity and sensitivity of this panel illustrates the potential of kinase activity as a predictive biomarker for treatment success of HER2-targeted therapy.

## Results

In order to identify kinase activities predictive of treatment success, we performed Quantakinome™ analysis on 32 treatment-naïve breast cancer biopsies. These clinical samples originated from the TRAIN-2 randomized phase 3 clinical trial (Van Der Voort et al. 2021; van Ramshorst et al. 2018) in which treatment-naive patients with stage II–III HER2+ breast cancer tumours were enrolled. All patients received therapy consisting of dual HER2 blockade (Trastuzumab and Pertuzumab) in combination with varying types of chemotherapy (with or without anthracyclines). The clinical outcome was defined as pathological complete response (pCR, no tumour cells left), near pathological complete response (npCR; <10% of cells contained invasive tumour cells) and no pathological complete response (No pCR; >10% tumour cells remaining). Patients classified as No pCR are referred to as treatment resistant throughout this study. Prior to drug treatment, a core needle biopsy was taken for local histology-based assessment and the remainder was stored for later analysis. In the current study, we received 32 of these pre-treatment tumour biopsy samples for QuantaKinome™ analysis. In total, 17 patients were classified as pCR, 6 as npCR and 9 as No pCR. A complete list of patient biopsy details is provided in Supplementary Table 1.

The tumour biopsies were lysed, and the proteins were extracted and digested before being desalted on an automated platform, and spiked with an internal reference standard containing stable-isotope labelled standards for each phosphorylated kinase T-loop. Subsequently, phosphorylated peptides were enriched by automated phosphopeptide enrichment. Thereafter, the signal of 311 T-loop peptides was measured using a single targeted mass spectrometry assay on a triple quadrupole mass spectrometer. Finally, endogenous kinase T-loop signals in each sample were normalised using their respective internal standard, to achieve a precise quantification of kinase activation. The complete workflow is depicted in Figure 1A.

**Figure 1.**
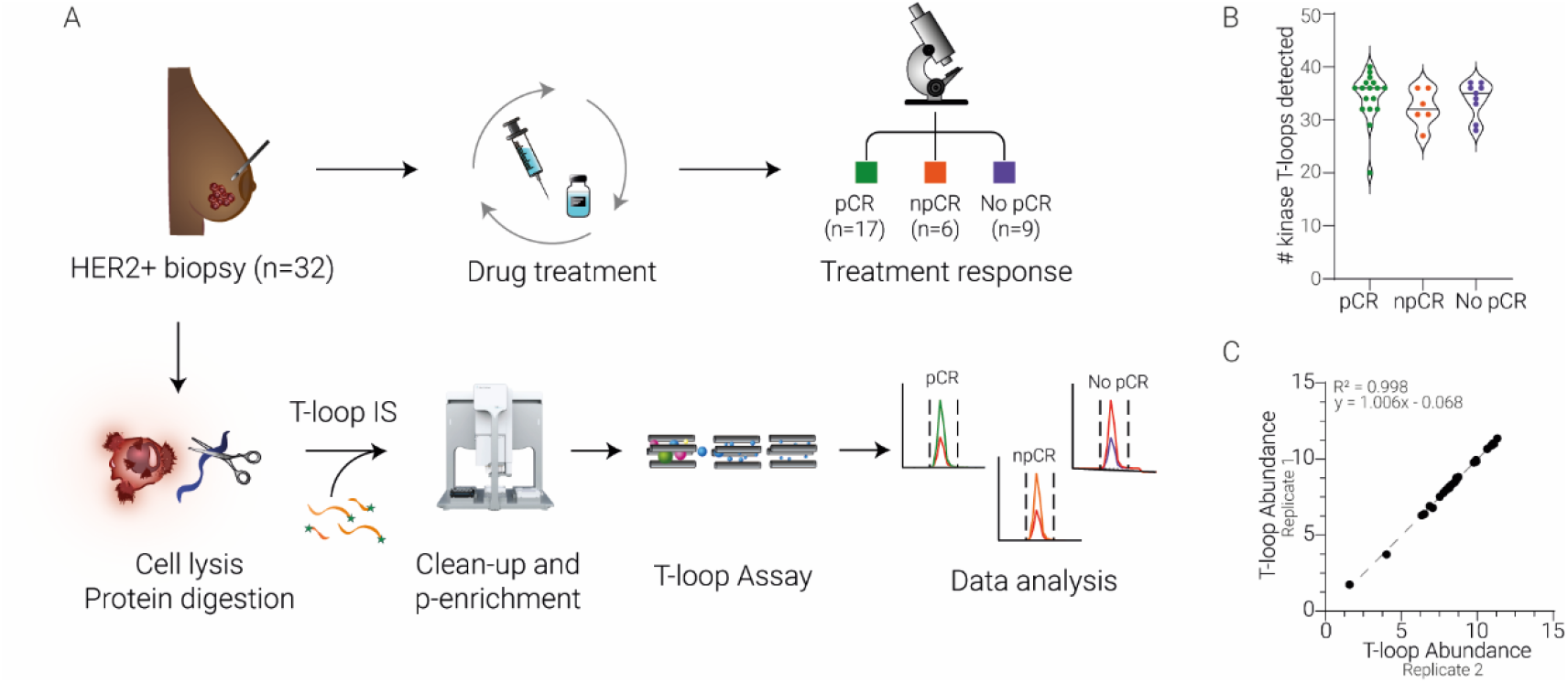
Study overview. A) Experimental workflow. B) Number of kinase T-loops quantified per patient biopsy in each treatment outcome group. C) Reproducibility of a technical workflow replicate

In total, we successfully detected 307 kinase T-loop reference standards of which 56 were endogenously quantified. 5 of the endogenously detected T-loop phosphopeptides contained a methionine oxidation and 4 were doubly phosphorylated, yielding a total of 51 unique kinase activation states, which were mapped to 61 endogenous kinases (Supplementary Figure 1A). The number of quantified kinase T-loops per patient ranged from 20 to 40 and was comparable between the three treatment outcome groups (Figure 1B).

One patient biopsy sample contained enough material to perform the entire workflow in duplicate on the exact same sample allowing us to explore the reproducibility of the workflow. As shown in Figure 1C, the correlation between the two replicates is very strong (R=0.998). The high correlation coefficient together with a slope of one and an offset near zero indicates a high reproducibility of the method in clinical samples.

A high dynamic range in the detection of kinases was found; the most abundant kinase T-loop (GSK3A [Y279]) was more than 4,400-fold higher abundant compared to the lowest detected kinase T-loop (NEK6 [S206]). Supplementary Figure 1B illustrates that all kinase families show a wide dynamic range and that the top five most abundant kinase T-loops are from the CMGC-family. Moreover, some kinases within the CMGC-family show high correlation, indicating that these kinases could be co-regulated. However, as shown in Supplementary Figure 1C, overall the kinase activity does not seem to be regulated family-wide, but rather on the individual kinase level. Typically, T-loop phosphorylation sites within the same kinase (such as CDK7 and MARK) have a high correlation.

The biological relevance of the detected kinases is evident from their high mutation frequency in breast cancer, which is similar to other well-known non-kinase breast cancer markers such as BRCA2 and ESR1 (Supplementary Figure 1D). Furthermore, Supplementary Figure 1E illustrates that the quantified kinase T-loops in this study consisted of both extensively studied kinases (such as ERK1/2, p38A and JNK1/2/3) and under-studied kinases.

### Low kinase activity in HER2-pathway is associated with treatment resistance

Since the targeted therapy used in this study is directed against HER2, we first explored the activity of kinases (by T-loop phosphorylation) relevant in this pathway. Firstly, the HER2-pathway was extensively covered in our dataset, including crucial kinases such as ERK, PDK1 and RSK1 (Figure 2A). Importantly, we found a large number of kinases in this pathway differentially regulated between the tumours that were treatment resistant (no pCR) compared to the treatment responsive tumours (pCR and npCR). The majority of differentially regulated kinases showed a lower kinase activity in treatment resistant tumours (such as ERK1, ERK2, RSK1, CaMKID and p38). However, one kinase was found upregulated amongst the resistant tumours: Focal Adhesion Kinase (FAK).

**Figure 2.**
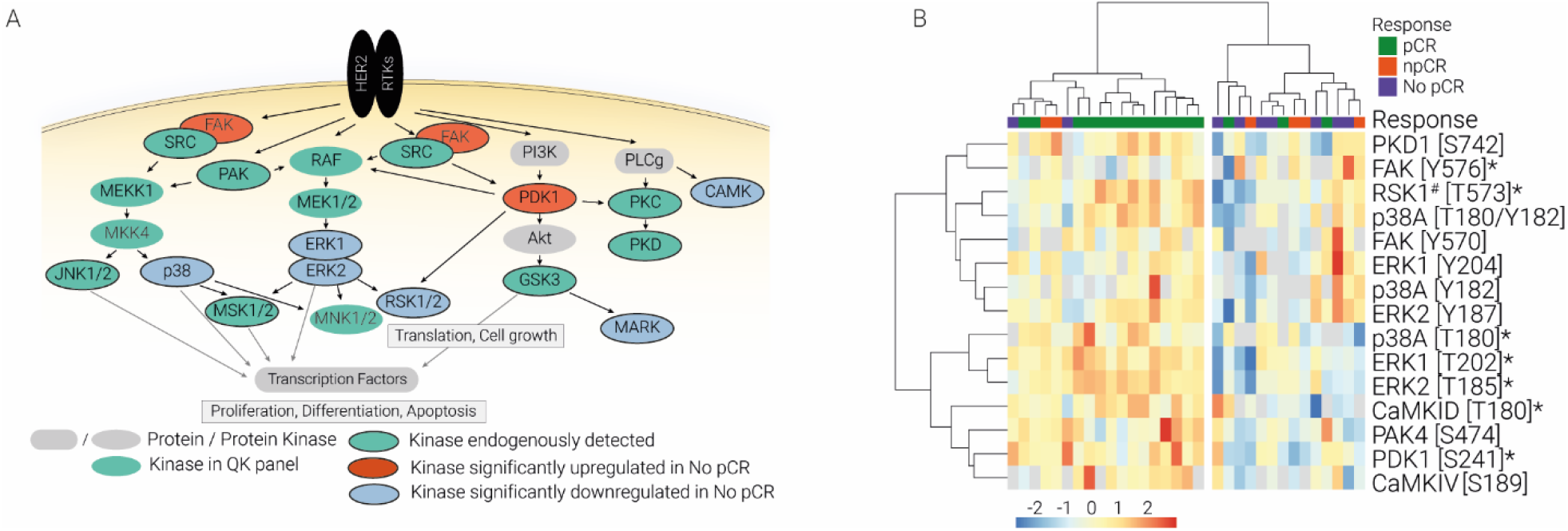
Low kinase activity levels within HER2 pathway are associated with treatment resistance. A) Graphical representation of HER2-pathway. Kinases quantified in this study are highlighted. B) Unsupervised clustering of HER2-pathway kinase activities (Euclidean distance, Z-scored data). ^*^Significantly regulated kinase; ^#^kinase T-loop peptide was shared with at least one paralog.

Unsupervised clustering of key kinases within the HER2-pathway grouped patients into two main clusters. One cluster contained the majority of the treatment resistant tumours and displayed low kinase activity levels (Figure 2B). Interestingly, we observed a kinase activity heterogeneity within patient groups, indicating that individual patients can have different nodes activated within the HER2-pathway, leading to a similar outcome. Overall, there is consensus on the pathway level: reduced kinase activity within the HER2-pathway was associated with treatment resistance. This could indicate that these tumours were less HER2-driven and hence less susceptible to HER2-inhibition.

### Differentially regulated kinase activities are predictive of treatment outcome

The primary goal of this study was to identify kinases that predict therapy response in HER2+ breast cancer treated with a combination of HER2-blockade and chemotherapy. An ANOVA-test revealed eleven kinases that were differentially activated between the pCR, npCR and No pCR groups. Unsupervised hierarchical clustering of these regulated kinases revealed two clusters of patients (Figure 3A). Cluster 1 consisted of mainly pCR tumours, which displayed a high relative kinase activity. Cluster 2 contained npCR and all No pCR tumours, characterised by generally lower kinase activity levels. As discusses above, many of these kinases were involved in the HER2-pathway.

**Figure 3.**
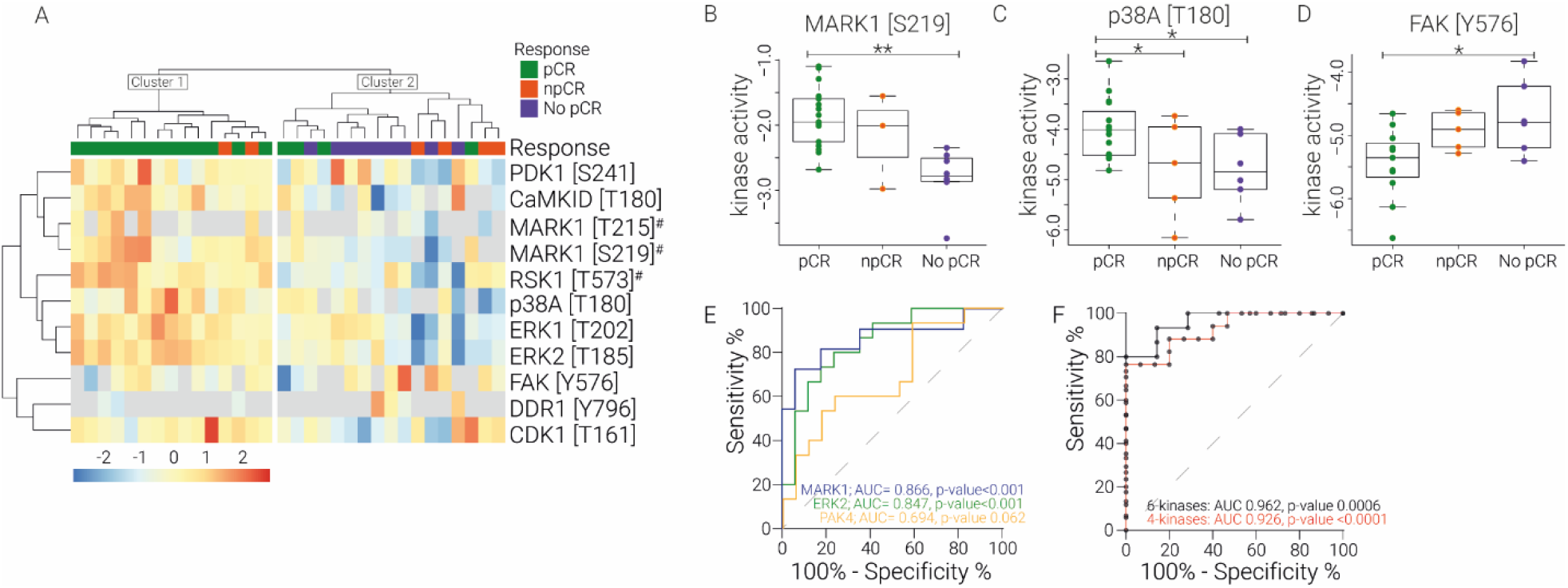
Kinase activities (by T-loop phosphorylation) differentiate treatment outcome groups. A) Heatmap of unsupervised clustering of all significantly regulated kinase T-loops (ANOVA p-value < 0.05) (Euclidean distance, data was Z-scored). ^#^ kinase T-loop was shared with at least one close paralog. B-D) Boxplots of regulated kinase T-loops. ^*^ p-value < 0.05; ^**^ p-value < 0.001. E) Receiver operator characteristics (ROC) curve analysis to predict treatment outcome based on kinase T-loop abundance for single kinases. F) ROC curve to predict treatment outcome based on a panel of kinases. Six-kinase panel consisted of kinase T-loops detected in at least 80% of the patients, with a minimum of 1.5 fold difference and max p-value of Four-kinase panel consisted of kinase T-loops detected in all patients with p-value < 0.05.

MARK (microtubule affinity-regulated kinase, also known as Par-1), RSK1, CDK1 and CaMKID were significantly downregulated amongst treatment resistant tumours and showed intermediate levels for the npCR tumours (Figure 3B, Supplementary Figure 2A). P38A, ERK1 and ERK2 contrarily were downregulated in both the npCR and No pCR tumours compared to the pCR tumours (Figure 3C, Supplementary Figure 2A). This suggests that p38A, ERK1 and ERK2 could be important in treatment sensitivity yet are not sufficient for a full treatment response, whereas MARK, RSK, CDK1 and CaMKID are stronger determinants for a complete treatment response. Apart from CDK1 and MARK, all of these kinases were involved in the HER2-pathway highlighting that reduced HER2-pathway kinase activity was linked to treatment resistance.

Interestingly, FAK showed an opposite trend, displaying a higher activity in the No pCR tumours (Figure 3D). FAK integrates signalling events from two different types of receptors; it is a key downstream kinase of both growth factor and integrin receptors. Hence, increased FAK activity suggests that these alternative signalling routes might be important in treatment resistant tumours and that tumours could escape HER2-inhibition via this alternative signalling node.

To assess whether kinase activity levels could be used to predict treatment outcome, a receiver-operating characteristic (ROC) curve analysis was performed using both single kinase activity levels and a panel of kinases. For clinically relevant predictions two patient groups were formed; pCR patients were considered “responders”, whereas No and npCR patients were designated “non-responders”. The kinase activity differences between these two groups and corresponding ROC characteristics are tabulated in Table I.

**Table I:**
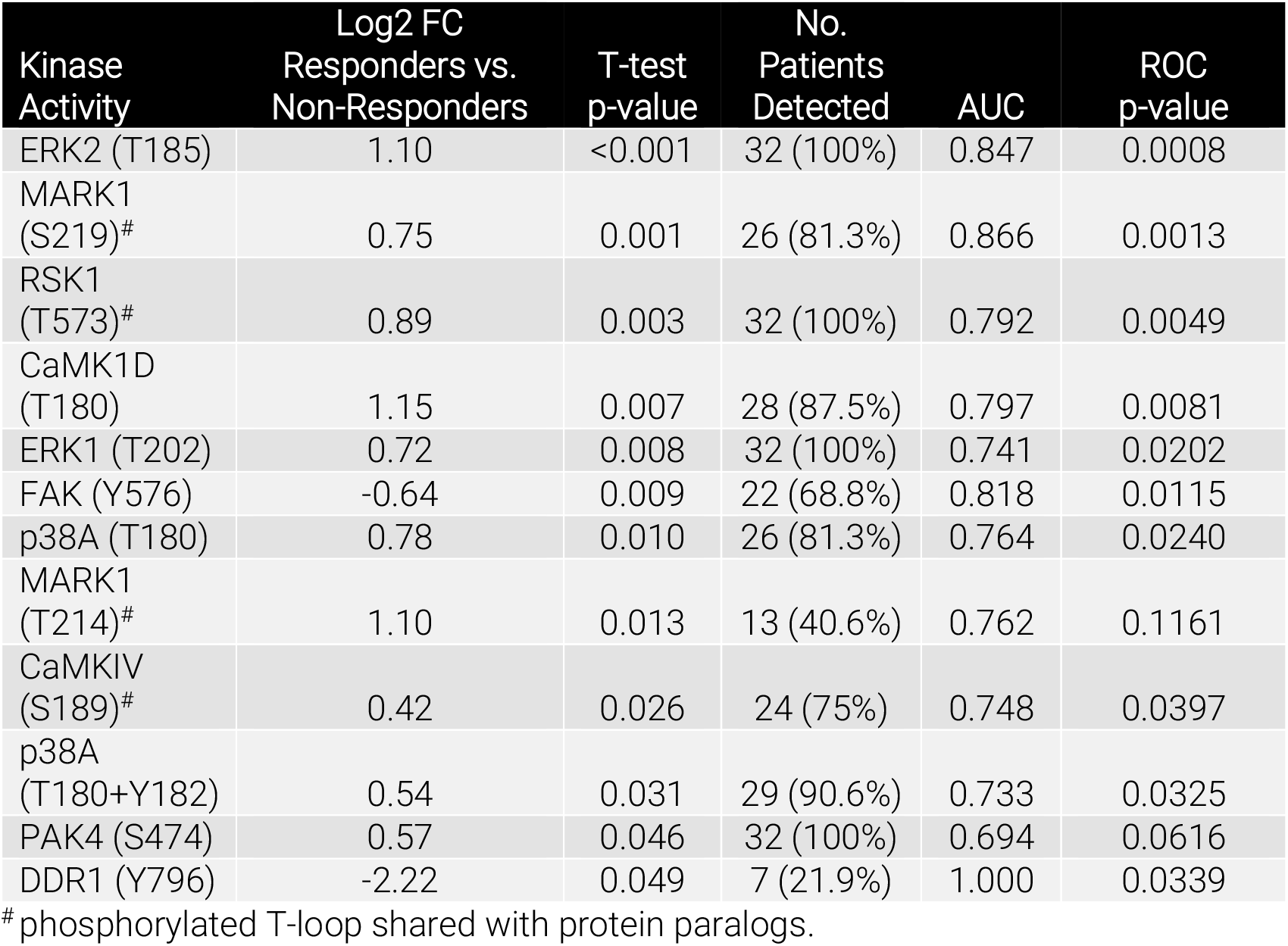
Kinase activities regulated in clinical groups and used for ROC analysis

The overall performance of our diagnostic markers to predict treatment outcome was assessed using the area under the curve (AUC) of the ROC. The AUC for the individual kinase activity measurements ranged from 0.866 to 0.694 (Table I, Figure 3E). The two most significantly regulated kinase activities (ERK2 T185 and MARK1 S219) showed good predictive potential with an AUC ≥0.85.

Previous research has shown that tumours display considerable heterogeneity between patients; tumours utilise different signalling routes to drive tumour growth or treatment evasion (Satpathy et al. 2020; Krug et al. 2020). Indeed, we also observed varying kinase activity levels between and within the patient groups in this study (Figure 2A and Figure 3A). Therefore, to account for this patient heterogeneity in a diagnostic test, we explored the use of a panel of kinase activities to better predict treatment response. The first kinase panel consisted of 6 kinase activities (RSK1 T573, CaMK1D T180, p38A T180, ERK1 T202, ERK2 T185, MARK1 S219) from the HER2-signalling pathway that were selected based on their p-value (<0.05), fold change (>1.5-fold) and detectability (detected in at least 80% of patients). For the second panel, kinase activation states were only considered if they were significantly regulated and detected in all patients. This resulted in a set of 4 kinases (RSK1 T573, ERK1 T202, ERK T185, PAK4 S474). A multiple logistic regression of both kinase signatures resulted in higher AUC values (0.962 and 0.926, respectively) compared to the best single kinase predictors (Figure 3F). In summary, the 4-kinase panel showed a strong basis for a prognostic test that performed better than the best single kinase T-loop (MARK1 S219) with an added benefit of 100% detectability compared to 81.3%.

## Conclusion and discussion

The significance of phosphorylation dynamics and kinase activity in health and disease is indisputable. In this study, we applied a method previously developed by Schmidlin et al. (Schmidlin et al. 2019) for the direct measurement of kinase activity (as measured by kinase T-loop phosphorylation) in 32 breast cancer biopsies. Results of this novel technology demonstrated a high reproducibility and sampling depth from low sample quantities. We quantified 56 kinase activation states, largely covering the HER2-pathway. Classical approaches use substrate phosphorylation as proxy for kinase activation; however, determination of the exact activated kinase is hindered by information-bias and overlapping substrate specificity. Our kinase T-loop assay contrarily, directly infers kinase activity from T-loop phosphorylation.

The primary goal of this study was to explore the potential of kinase activation states in predicting therapy response in HER2+ breast cancer patients. 11 kinase activation states were significantly regulated between the different response groups. Patients that responded well to therapy showed a generally higher kinase activation profile, especially in the HER2-pathway. Poor responders displayed reduced kinase activities within the HER2-pathway, indicating a lower HER2-dependency among these tumours and hence poor response to HER2-inhibition. This is in line with previous research showing that decreased HER2-expression levels were linked to unfavourable treatment outcomes (Nuciforo et al. 2016; Baselga et al. 2014). Importantly, we observed heterogeneity on the individual kinase level within the HER2 pathway, highlighting the importance of a broad coverage of signalling nodes to catch tumour heterogeneity for mapping pathway activation.

The prognostic value of kinase activation levels was demonstrated by ROC analysis. Several kinase activation states were indeed predictive for treatment success of HER2-targeted therapy. Furthermore, the prognostic power was improved when a panel of kinase activities was used; the benefit of this panel compared to single kinase activities likely results from the heterogeneity in activating signalling nodes between patients. Hence, combining the activation states of multiple kinases within a single diagnostic test is crucial.

In this study, we found significant downregulation of p38A [T180] amongst the treatment resistant tumours. Previous research has revealed the highly complex and context-specific function of p38 in drug resistance in cancer. In leukaemia, upregulation of p38 is linked to drug resistance against genotoxic chemotherapy (Gao and Liu 2016), whereas the opposite holds true for targeted therapy; increased p38 phosphorylation is then linked to increased drug sensitivity (Parmar et al. 2004; Dumka et al. 2009). Furthermore, resistance against EGFR inhibitors in lung cancer can be overcome by dual inhibition of MEK and PI3K via activation of p38 signalling (Sato et al. 2018). The antibody-based techniques used as readout for p38-activation could partially explain the different roles attributed to this kinase, since distinct biological functions have been ascribed to different phosphorylation states of p38 (Mittelstadt et al. 2009). Due to the close proximity of these phosphorylation sites in the kinase T-loop, these are difficult to distinguish using antibodies. In our study, we quantified multiple T-loop activation states of p38A, of which only p38A [T180] was found significantly changing (Supplementary Figure 2B). This suggests diverse functions of p38A activation states and highlights the potential of our method to distinguish between these kinase states.

Among the treatment resistant tumours, we found a significant upregulation of FAK activity at FAK Y576. A growing body of evidence suggests that FAK may play an important role in cancer biology and therapy resistance and a variety of FAK-inhibitors is currently in development (Lee et al. 2015; Lv et al. 2018). FAK has been found over-expressed and/or hyper-phosphorylated in many cancers, including breast cancer (Weiner, Liu, and Craven 1990; Park et al. 2010). This over-expression or over-activation is associated with increased cell motility, survival, proliferation and poor clinical outcome (Balsas et al. 2017; Ferguson and Gray 2018; Pylayeva et al. 2009; Fan et al. 2016). Furthermore, FAK has been linked to Trastuzumab resistance via compensatory cross-talk (Park et al. 2010) and inhibition of FAK has been shown to help to overcome this resistance (Lazaro et al. 2013). Our data is in agreement with these findings and suggests that over-activation of FAK might indeed play a role in reduced drug sensitivity. The increased FAK activity could provide alternative signalling nodes aiding in the tumour’s escape of HER2-inhibition.

In addition to the differential regulation of these well-described kinases, we also identified a significant downregulation of the relatively unexplored kinase MARK among treatment resistant tumours. This serine-threonine kinase is often found amplified in breast cancer (Supplementary Figure 1D) and plays a role in cell motility and regulation of energy metabolism. Recently, MARK1 has been discovered as the direct target of microRNA in both cervical and colorectal cancer where it was linked to proliferation and cell migration (Natalia et al. 2018; Tang et al. 2019). Furthermore, MARK is a substrate of LKB1, a well-known tumour suppressor gene linked to metastatic outgrowth of cancer cells (Lizcano et al. 2004; Spicer et al. 2003; Goodwin et al. 2014). In this study, we link increased MARK activation to better treatment response and show that the activation status of this kinase is predictive for drug resistance in this cohort of patients. Since computational tools used in shotgun phosphoproteomics exploit substrate phosphorylation as surrogate for kinase activity, it is inherently biased towards well-studied kinases. Hence, a kinase such as MARK (for which the downstream targets are largely unknown) will remain hidden in these conventional methods. The direct measurement of kinase activity used in this study does not suffer from this information-bias and is therefore able to identify this kinase as a potential new player in drug resistance.

The suitability of the kinase T-loop assay as a diagnostic tool in breast cancer treatment requires further validation in larger clinical cohorts. However, we believe that our study shows the large potential of this technology in the prediction of treatment response *in vivo*.

## Acknowledgments

DOD, KES and MA acknowledge support from the Horizon 2020 program INFRAIA project Epic-XS (Project 823839) and the NWO funded Netherlands Proteomics Centre through the National Road Map for Large-scale Infrastructures program X-Omics (Project 184.034.019) of the Netherlands Proteomics Centre.

We would like to thank Lennart Mulder for technical assistance and the NKI-AVL Core Facility Molecular Pathology & Biobanking (CFMPB) for supplying NKI-AVL Biobank material.

## Method

### KEY RECOURCES TABLE

KEY REOURCES TABLE
| REAGENT | SOURCE | IDENTIFIER |
| --- | --- | --- |
| **Biological samples** |  |  |
| Human IBC tumour samples | TRAIN2 clinical trial | https://clinicaltrials.gov/ct2/show/NCT01996267 |
| **Chemicals, peptides and recombinant proteins** |  |  |
| Sodium deoxycholate (SDC) | Sigma Aldrich | MFCD00064139 |
| Tris(2- carboxyethyl) phosphinehydrochloride (TCEP) | Sigma Aldrich | MFCD00145469 |
| Chloroacetamide (CAA) | Sigma Aldrich | MFCD0008027 |
| TRIS | Sigma Aldrich |  |
| PhosSTOP | Merck | 4906837001 |
| cOmplete, Mini, EDTA-free Protease Inhibitor Cocktail | Merck | 11836170001 |
| Trypsin | Thermo Scientific | Prod# 90057S |
| LysC | Wako | 125-05061 |
| SpikeMix Kinase Activation Loops (Human) - heavy | JPT Germany | SPT-KAL-POOL-L-100pm |
| iRT Kit | Biognosys | Ki-3002-1 |
| **Critical commercial assays** |  |  |
| AssayMap Cartridge Rack, Fe(III)-NTA 5 mL | Agilent Technologies | Cat#G5496-60085 |
| AssayMAP Cartridge Rack C18 5ul | Agilent Technologies | Cat# 5190-6532 |
| Bradford Protein Assay | Bio-Rad | 5000006 |
| PepMap RSLC C18 2um, 100A 75x25 | Thermo Scientific | ES802 |
| **Deposited Data** |  |  |
| Skyline documents | Panorama |  |
| Raw files | Panorama |  |
| **Software and algorithms** |  |  |
| Skyline | Skyline | https://skyline.ms/project/home/software/Skyline/ |
| GraphPad Prism 9.3.0 | Graphpad Software Inc | https://www.graphpad.com/scientific-software/prism/ |
| R v4.3.3 | https://www.r-project.org/ | https://www.r-project.org/ |
| RStudio (v2022.02.1 461) | RStudio, PBC | https://www.rstudio.com/ |

### RESOURCE AVAILIBILITY

#### Lead contact

Further information and requests for resources and reagents should be directed to and will be fulfilled by the lead contact, Maarten Altelaar.

#### Materials availability

The study did not generate new unique reagents.

#### Data and code availability

- The mass spectrometry data used in this study has been deposited to Panorama
- This study did not generate original code
- Any additional information required to reanalyses the data reported in this paper is available form the lead contact upon request

### EXPERIMENTAL MODEL AND SUBJECT DETAILS

Patient biopsies were obtained from patients enrolled in the TRAIN-2 study (van Ramshorst et al., 2016).

## METHOD DETAILS

### Biopsy preparation

Patient biopsies were obtained from patients enrolled in the TRAIN-2 study (van Ramshorst et al. 2016). The study was approved by the ethical committee and informed consent was obtained. Patients overexpressed HER2 and received neoadjuvant therapy consisting of Trastuzumab and Pertuzumab supplemented with either 5-fluorouracil, epirubicin, cyclophosphamide or a combination of paclitaxel and carboplatin. Prior to start of the treatment a 14G needle biopsy was taken of approximately 30 μm and flash frozen in liquid nitrogen. Part of this biopsy was used to perform hematoxylin and eosin (HE) staining to determine tumour cell content, while 1/3 of the biopsy was snap frozen and kept at -80°C for QuantaKinome™ analysis. Only samples with a minimum tumour percentage of 60% were used in this study. After nine cycles of drug treatment, the response was determined at surgery.

### Kinase activity analysis using QuantaKinome™

The QuantaKinome™ platform was applied to measure T-loop phosphopeptides by using a targeted LC-MS approach (QuantaKinome™, Pepscope B.V.), which is based on the method described in Schmidlin et al. Briefly, frozen biopsy were lysed and sonicated in lysis buffer. For each sample, 200 μg of protein was processed. After reduction, alkylation and digestion, all samples were dried and stored at -20°C until phosphoenrichment. Phosphorylated peptides were desalted and enriched using an automated platform. Samples were dried and stored at -80°C until LC-MS analysis. Next, samples within one experiment were measured in randomized order using the QuantaKinome™ targeted LC-MS assay (QuantaKinome™ Library v1, Pepscope).

## QUANTIFICATION AND STATISTICAL ANALYSIS

### Preprocessing of datasets

Raw files were uploaded into Skyline (MacLean et al. 2010) and peak boundaries were manually checked and adjusted if needed. Subsequently, an in-house build R-script was used to filter-out endogenous peptides with interference. First, all transitions with a signal-to-noise level below three times the background were removed. Subsequently, for each peptide the relative contribution of each transition to the total peptide signal was calculated and light transitions with a difference of more than 20% compared to the heavy standard relative contribution were removed. Next, peptides with less than two transitions were removed. Lastly, peptides that were detected in less than five files were removed. Remaining peptides were visually inspected in Skyline. After peptide identification, the transitions suitable for quantification were chosen to reliably quantify the peptides over all samples. For each peptide, only transitions that showed a consistent light/heavy ratio in all samples were used. Furthermore, quantification was based on at least two transitions. The same transitions were used to quantify across all files. Lastly, a weighted and internal standard corrected T-loop abundance was created by dividing the sum of the light transitions by the sum of the heavy transitions following a log2 transformation. The T-loop abundance is also referred to as kinase activity in this manuscript.

Some peptides were detected in oxidised and non-oxidised form. Oxidised T-loop phosphopeptides were detected less frequently and generally showed a 10-fold lower signal compared to their non-oxidised counterpart (Supplementary Figure 3). The correlation between the two oxidation forms was high in all samples, indicating that the oxidation rate was similar between all samples. Therefore, when both oxidation states were detected, only the non-oxidised form was used to calculate the kinase T-loop abundance level.

### Statistics

To compare three means, a one-way ANOVA test was used (using the aov-function, followed by TukeyHSD-function in Rstudio). A p-value below 0.05 was regarded as statistically significant. Correlation analysis was performed by Pearson correlation using cor-function in RStudio.

Unsupervised clustering and visualization were performed in R using the pheatmap package. The data was z-scored (scaled by row) prior to clustering. Euclidean distance was used for both row and column clustering. ROC analysis and multiple logistic regression was performed in Graphpad Prism 9.

### Data and code availability

Raw data and processed data are available on Panorama.

---

**Supplementary Table 1.** Patient information

| *Patient* | *Histology* | *Tumour grade* | *ER (%)* | *PR (%)* | *HER2 score* | *Treatment outcome* | *Tumour (%)* |
| --- | --- | --- | --- | --- | --- | --- | --- |
| 1 | ID | 2 | 95 | 70 | 3+ | pCR | 80 |
| 2 | ID | 2 | 100 | 0 | 3+ | pCR | 80 |
| 3 | ID | 3 | 20 | 0 | 3+ | pCR | 80 |
| 4 | ID | 2 | 80 | 5 | 3+ | pCR | 70 |
| 5 | IL | 3 | 100 | 40 | 3+ | npCR | 80 |
| 6 | ID | 2 | 40 | 0 | 3+ | pCR | 60 |
| 7 | ID | 2 | 100 | 1 | 2+ | pCR | 70 |
| 9 | ID | 3 | 80 | 70 | 3+ | npCR | 70 |
| 10 | ID | 2 | 0 | 0 | 3+ | pCR | 70 |
| 11A* | ID | 2 | 0 | 0 | 3+ | pCR | 60 |
| 11B* | ID | 2 | 0 | 0 | 3+ | pCR | 60 |
| 12 | ID | 3 | 0 | 0 | 3+ | pCR | 70 |
| 13 | ID | 2 | 0 | 0 | 3+ | pCR | 60 |
| 14 | ID | 3 | 60 | 5 | 2+ | npCR | 60 |
| 15 | ID | 3 | 0 | 0 | 3+ | pCR | 60 |
| 16 | IL | 3 | 90 | 0 | 3+ | pCR | 80 |
| 17 | ID | 3 | 90 | 100 | 3+ | npCR | 70 |
| 18 | ID | 3 | 100 | 20 | 3+ | No pCR | 80 |
| 20 | ID | 3 | 90 | 40 | 3+ | pCR | 60 |
| 21 | ID | 3 | 99 | 1 | 2+ | No pCR | 80 |
| 22 | ID | 3 | 100 | 60 | 2+ | No pCR | 80 |
| 23 | ID | 2 | 70 | 70 | 3+ | pCR | 60 |
| 24 | ID | 3 | 10 | 0 | 2+ | pCR | 80 |
| 25 | ID | 3 | 100 | 100 | 3+ | pCR | 70 |
| 26 | ID | 2 | 50 | 100 | 3+ | No pCR | 70 |
| 27 | ID | 3 | 100 | 5 | 3+ | No pCR | 80 |
| 28 | ID | 3 | 100 | 90 | 3+ | No pCR | 70 |
| 30 | ID | 2 | 100 | 40 | 3+ | No pCR | 60 |
| 32 | ID | 3 | 100 | 1 | 3+ | npCR | 60 |
| 33 | ID | 3 | 100 | 0 | 2+ | pCR | 80 |
| 34 | ID | 2 | 100 | 0 | 3+ | npCR | 70 |
| 36 | ID | 3 | 100 | 5 | 3+ | No pCR | 80 |
| 37 | IL | 2 | 100 | 0 | 2+ | No pCR | 80 |

ID: invasive ductal
IL: invasive lobular
\*Workflow replicate

## Supplementary Figures

**Supplementary Figure 1.**
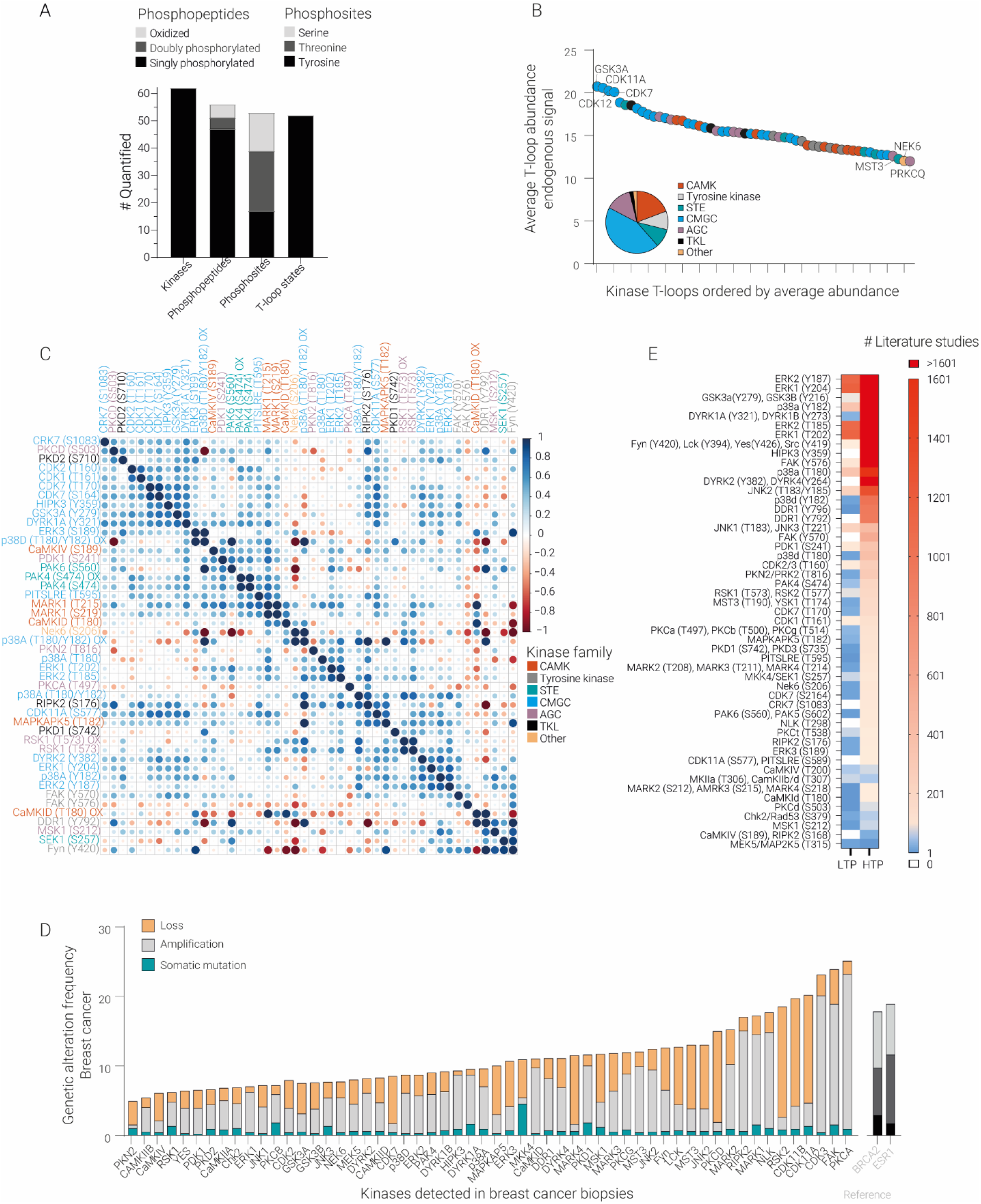
Characterisation of quantified T-loops and kinases in this study. A) Total number of quantified kinases, phosphopeptides, phosphosites and T-loops across all biopsies. B) Average kinase T-loop abundance of the endogenous signals, coloured by kinase family. Amongst the most highly abundant kinases are family members of the CMGC-family (GSK3A and CDKs). C) Correlation plot of all quantified kinase T-loops in this study. Kinase names are coloured according to the kinase family. Pearson correlation was used. Hierarchical clustering method was Ward.D. D) Kinase mutation frequency in breast cancer, reported by TCGA (The Cancer Genome Atlas). Many detected kinases show a very high mutation frequency in breast cancer, comparable to other well-known breast cancer associated genes such as BRCA and ESR1. E) Number of references in PhosphoSitePlus for each endogenously detected T-loop phosphosite. Although many (but not all) sites have been identified often in high throughput studies (HTS), the biological relevance has been mostly understudied, as evidenced by the low number of low througphut studies (LTS).

**Supplementary Figure 2.**
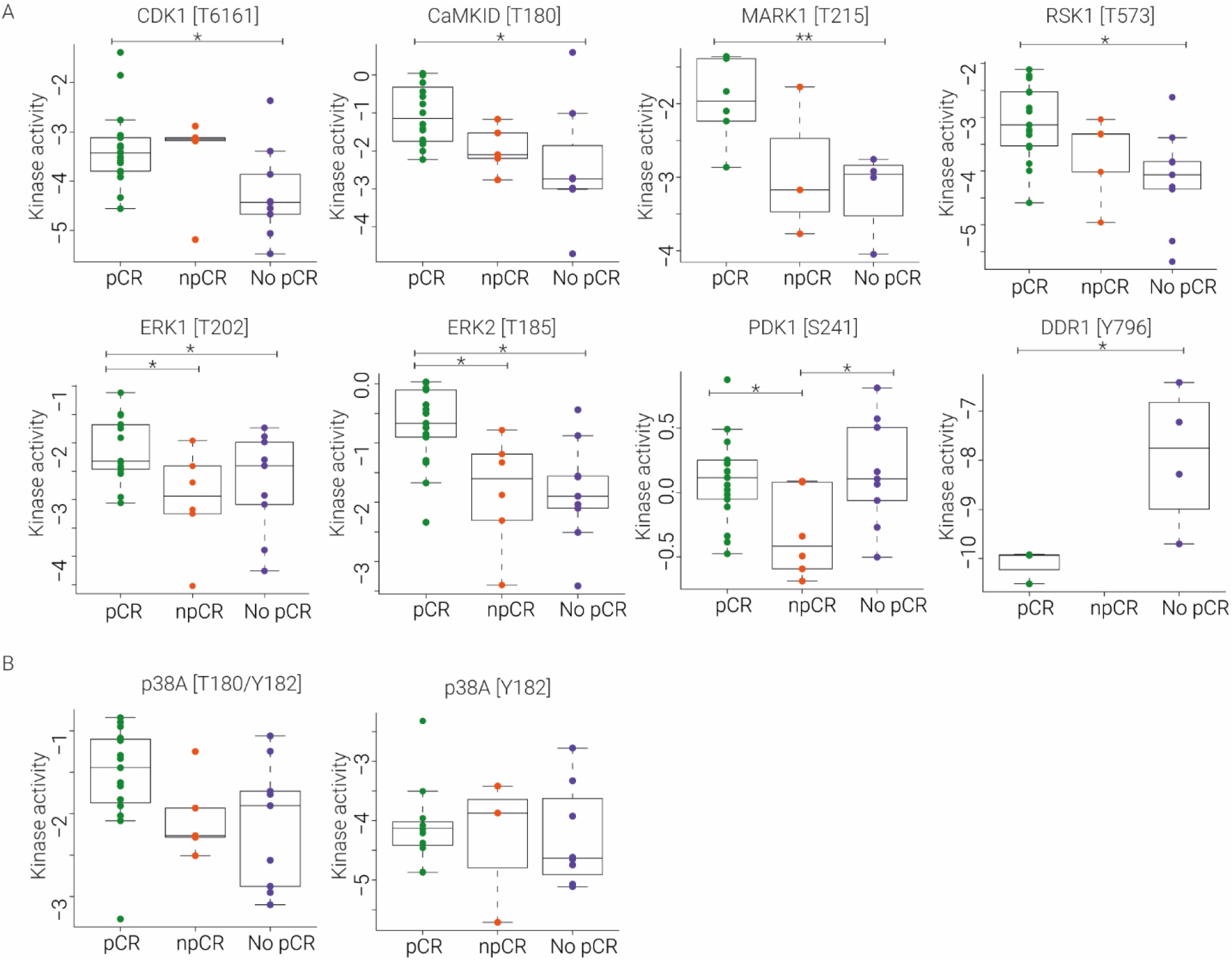
Boxplots of kinase T-loops. A) Boxplots of differentially regulated kinase T-loops. ^*^ p-value < 0.05; ^**^ p-value < 0.001. B) Boxplots of kinase T-loops of p38A that were not significantly changing between treatment outcome groups.

**Supplementary Figure 3.**
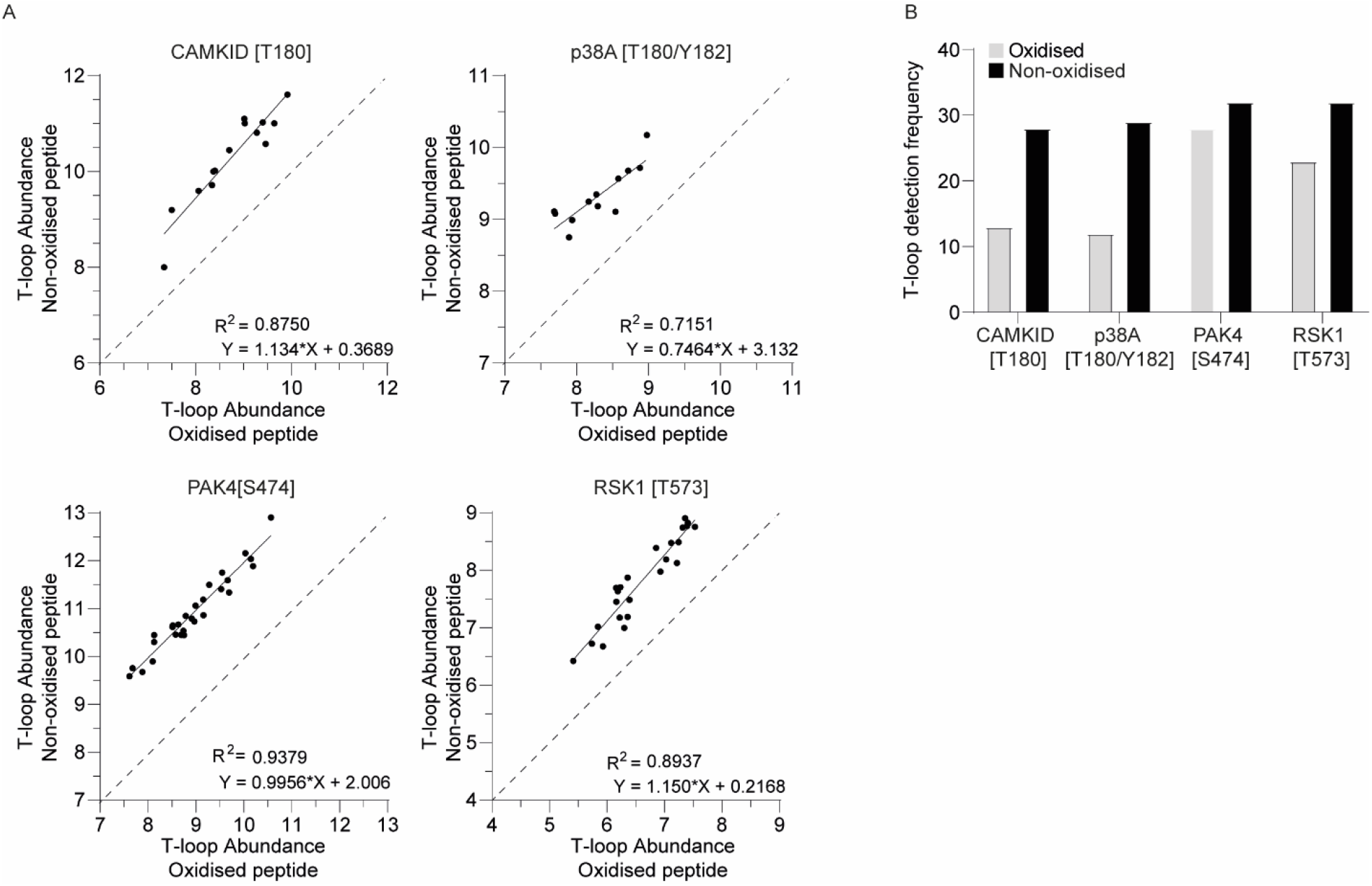
Detected oxidised and non-oxidised kinase T-loops. A) Correlation plots, comparing the abundance of the oxidised and non-oxidised T-loop versions. Good linear correlations show the rate of oxidation is similar between samples. Moreover, oxidised versions are >10-fold lower in abundance. B) The number of quantified T loops is lower for the oxidised T loop phosphopeptides.

## References

Andrulis, Irene L., Shelley B. Bull, Martin E. Blackstein, Donald Sutherland, Carmen Mak, Saul Sidlofsky, Kenneth P.H. Pritzker, et al. 1998. “Neu/ErbB-2 Amplification Identifies a Poor-Prognosis Group of Women with Node-Negative Breast Cancer.” Journal of Clinical Oncology 16 (4): 1340–49. https://doi.org/10.1200/JCO.1998.16.4.1340.

Balsas, Patricia, Jara Palomero, Álvaro Eguileor, Marta Leonor Rodríguez, Maria Carmela Vegliante, Ester Planas-Rigol, Marta Sureda-Gómez, Maria C. Cid, Elias Campo, and Virginia Amador. 2017. “SOX11 Promotes Tumor Protective Microenvironment Interactions through CXCR4 and FAK Regulation in Mantle Cell Lymphoma.” Blood 130 (4): 501–13. https://doi.org/10.1182/blood-2017-04-776740.

Baselga, José, Javier Cortés, Seock-Ah Im, Emma Clark, Graham Ross, Astrid Kiermaier, and Sandra M Swain. 2014. “Biomarker Analyses in CLEOPATRA: A Phase III, Placebo-Controlled Study of Pertuzumab in Human Epidermal Growth Factor Receptor 2-Positive, First-Line Metastatic Breast Cancer.” Journal of Clinical Oncology : Official Journal of the American Society of Clinical Oncology 32 (33): 3753–61. https://doi.org/10.1200/JCO.2013.54.5384.

Canovas, Begoña, and Angel R. Nebreda. 2021. “Diversity and Versatility of P38 Kinase Signalling in Health and Disease.” Nature Reviews Molecular Cell Biology 22 (5): 346–66. https://doi.org/10.1038/s41580-020-00322-w.

Carter, Paul, Len Presta, Cornelia M. Gorman, John B.B. Ridgway, Dennis Henner, Wai Lee T. Wong, Ann M. Rowland, Claire Kotts, Monique E. Carver, and H. Michael Shepard. 1992. “Humanization of an Anti-P185HER2 Antibody for Human Cancer Therapy.” Proceedings of the National Academy of Sciences of the United States of America 89 (10): 4285–89. https://doi.org/10.1073/pnas.89.10.4285.

Cohen, P. 2001. “The Role of Protein Phosphorylation in Human Health and Disease. The Sir Hans Krebs Medal Lecture.” European Journal of Biochemistry. England. https://doi.org/10.1046/j.0014-2956.2001.02473.x.

Cohen, Philip. 2002. “The Origins of Protein Phosphorylation.” Nature Cell Biology 4 (5). https://doi.org/10.1038/ncb0502-e127.

Dumka, Disha, Poonam Puri, Nathalie Carayol, Crystal Lumby, Harikrishnan Balachandran, Katja Schuster, Amit K Verma, Lance S Terada, Leonidas C Platanias, and Simrit Parmar. 2009. “Activation of the P38 Map Kinase Pathway Is Essential for the Antileukemic Effects of Dasatinib.” Leukemia & Lymphoma 50 (12): 2017–29. https://doi.org/10.3109/10428190903147637.

Fan, Tianli, Jing Chen, Lirong Zhang, Pan Gao, Yiran Hui, Peirong Xu, Xiaqing Zhang, and Hongtao Liu. 2016. “Bit1 Knockdown Contributes to Growth Suppression as Well as the Decreases of Migration and Invasion Abilities in Esophageal Squamous Cell Carcinoma via Suppressing FAK-Paxillin Pathway.” Molecular Cancer 15 (1): 1–14. https://doi.org/10.1186/s12943-016-0507-5.

Ferguson, Fleur M., and Nathanael S. Gray. 2018. “Kinase Inhibitors: The Road Ahead.” Nature Reviews Drug Discovery 17 (5): 353–76. https://doi.org/10.1038/nrd.2018.21.

Gao, F, and W-J Liu. 2016. “Advance in the Study on P38 MAPK Mediated Drug Resistance in Leukemia.” European Review for Medical and Pharmacological Sciences 20 (6): 1064–70.

Goodwin, Jonathan M., Robert U. Svensson, Hua Jane Lou, Monte M. Winslow, Benjamin E. Turk, and Reuben J. Shaw. 2014. “An AMPK-Independent Signaling Pathway Downstream of the LKB1 Tumor Suppressor Controls Snail1 and Metastatic Potential.” Molecular Cell 55 (3): 436–50. https://doi.org/10.1016/j.molcel.2014.06.021.

Harbeck, Nadia, Frédérique Penault-Llorca, Javier Cortes, Michael Gnant, Nehmat Houssami, Philip Poortmans, Kathryn Ruddy, Janice Tsang, and Fatima Cardoso. 2019. Breast Cancer. Nature Reviews Disease Primers. Vol. 5. https://doi.org/10.1038/s41572-019-0111-2.

Hu, Zhiyuan, Cheng Fan, Daniel S. Oh, J. S. Marron, Xiaping He, Bahjat F. Qaqish, Chad Livasy, et al. 2006. “The Molecular Portraits of Breast Tumors Are Conserved across Microarray Platforms.” BMC Genomics 7: 1–12. https://doi.org/10.1186/1471-2164-7-96.

Krug, Karsten, Eric J Jaehnig, Shankha Satpathy, Lili Blumenberg, Alla Karpova, Meenakshi Anurag, George Miles, et al. 2020. “Proteogenomic Landscape of Breast Cancer Tumorigenesis and Targeted Therapy.” Cell 183 (5): 1436-1456.e31. https://doi.org/10.1016/j.cell.2020.10.036.

Lazaro, Glorianne, Chris Smith, Lindy Goddard, Nicola Jordan, Richard McClelland, Peter Barrett-Lee, Robert Nicholson, and Stephen Hiscox. 2013. “Targeting Focal Adhesion Kinase in ER+/HER2+ Breast Cancer Improves Trastuzumab Response.” Endocrine-Related Cancer 20 (5): 691–704. https://doi.org/10.1530/ERC-13-0019.

Lee, Brian Y., Paul Timpson, Lisa G. Horvath, and Roger J. Daly. 2015. “FAK Signaling in Human Cancer as a Target for Therapeutics.” Pharmacology & Therapeutics 146: 132–49. https://doi.org/10.1016/j.pharmthera.2014.10.001.

Lizcano, Jose M, Olga Göransson, Rachel Toth, Maria Deak, Nick A Morrice, Jérôme Boudeau, Simon A Hawley, et al. 2004. “LKB1 Is a Master Kinase That Activates 13 Kinases of the AMPK Subfamily, Including MARK/PAR-1.” The EMBO Journal 23 (4): 833–43. https://doi.org/10.1038/sj.emboj.7600110.

Lv, Peng Cheng, Ai Qin Jiang, Wei Ming Zhang, and Hai Liang Zhu. 2018. “FAK Inhibitors in Cancer, a Patent Review.” Expert Opinion on Therapeutic Patents 28 (2): 139–45. https://doi.org/10.1080/13543776.2018.1414183.

MacLean, Brendan, Daniela M. Tomazela, Nicholas Shulman, Matthew Chambers, Gregory L. Finney, Barbara Frewen, Randall Kern, David L. Tabb, Daniel C. Liebler, and Michael J. MacCoss. 2010. “Skyline: An Open Source Document Editor for Creating and Analyzing Targeted Proteomics Experiments.” Bioinformatics 26 (7): 966–68. https://doi.org/10.1093/bioinformatics/btq054.

Michailidou, Asimina, Hans-Jörg Trenz, and Pieter de Wilde. 2019. “Annex I:” The Internet and European Integration, 167–72. https://doi.org/10.2307/j.ctvdf0dxq.12.

Mittelstadt, Paul R., Hiroshi Yamaguchi, Ettore Appella, and Jonathan D. Ashwell. 2009. “T Cell Receptor-Mediated Activation of P38α by Mono-Phosphorylation of the Activation Loop Results in Altered Substrate Specificity.” Journal of Biological Chemistry 284 (23): 15469–74. https://doi.org/10.1074/jbc.M901004200.

Natalia, Martinez-Acuna, Gonzalez-Torres Alejandro, Tapia-Vieyra Juana Virginia, and Luis Marat Alvarez-Salas. 2018. “MARK1 Is a Novel Target for MiR-125a-5p: Implications for Cell Migration in Cervical Tumor Cells.” MicroRNA (Shariqah, United Arab Emirates) 7 (1): 54–61. https://doi.org/10.2174/2211536606666171024160244.

Nolen, Brad, Susan Taylor, and Gourisankar Ghosh. 2004. “Regulation of Protein Kinases: Controlling Activity through Activation Segment Conformation.” Molecular Cell 15 (5): 661–75. https://doi.org/10.1016/j.molcel.2004.08.024.

Nuciforo, Paolo, Sheeno Thyparambil, Claudia Aura, Ana Garrido-Castro, Marta Vilaro, Vicente Peg, José Jimenez, et al. 2016. “High HER2 Protein Levels Correlate with Increased Survival in Breast Cancer Patients Treated with Anti-HER2 Therapy.” Molecular Oncology 10 (1): 138–47. https://doi.org/10.1016/j.molonc.2015.09.002.

Owens, Marilyn A., Bruce C. Horten, and Moacyr M. Da Silva. 2004. “HER2 Amplification Ratios by Fluorescence in Situ Hybridization and Correlation with Immunohistochemistry in a Cohort of 6556 Breast Cancer Tissues.” Clinical Breast Cancer 5 (1): 63–69. https://doi.org/10.3816/CBC.2004.n.011.

Park, Joo Hyun, Byung Lan Lee, Jiyeon Yoon, Jin Kim, Min A. Kim, Han Kwang Yang, and Woo Ho Kim. 2010. “Focal Adhesion Kinase (FAK) Gene Amplification and Its Clinical Implications in Gastric Cancer.” Human Pathology 41 (12): 1664–73. https://doi.org/10.1016/j.humpath.2010.06.004.

Parmar, Simrit, Efstratios Katsoulidis, Amit Verma, Yongzhong Li, Antonella Sassano, Lakhvir Lal, Beata Majchrzak, et al. 2004. “Role of the P38 Mitogen-Activated Protein Kinase Pathway in the Generation of the Effects of Imatinib Mesylate (STI571) in BCR-ABL-Expressing Cells.” The Journal of Biological Chemistry 279 (24): 25345–52. https://doi.org/10.1074/jbc.M400590200.

Pernas, Sonia, and Sara M Tolaney. 2019. “HER2-Positive Breast Cancer: New Therapeutic Frontiers and Overcoming Resistance.” Therapeutic Advances in Medical Oncology 11: 1758835919833519. https://doi.org/10.1177/1758835919833519.

Pylayeva, Yuliya, Kelly M Gillen, William Gerald, Hilary E Beggs, Louis F Reichardt, and Filippo G Giancotti. 2009. “Ras- and PI3K-Dependent Breast Tumorigenesis in Mice and Humans Requires Focal Adhesion Kinase Signaling.” The Journal of Clinical Investigation 119 (2): 252–66. https://doi.org/10.1172/JCI37160.

Ramshorst, Mette S. van, Erik van Werkhoven, Aafke H. Honkoop, Vincent O. Dezentjé, Irma M. Oving, Ingrid A. Mandjes, Inge Kemper, et al. 2016. “Toxicity of Dual HER2-Blockade with Pertuzumab Added to Anthracycline versus Non-Anthracycline Containing Chemotherapy as Neoadjuvant Treatment in HER2-Positive Breast Cancer: The TRAIN-2 Study.” Breast 29: 153–59. https://doi.org/10.1016/j.breast.2016.07.017.

Ramshorst, Mette S van, Anna van der Voort, Erik D van Werkhoven, Ingrid A Mandjes, Inge Kemper, Vincent O Dezentjé, Irma M Oving, et al. 2018. “Neoadjuvant Chemotherapy with or without Anthracyclines in the Presence of Dual HER2 Blockade for HER2-Positive Breast Cancer (TRAIN-2): A Multicentre, Open-Label, Randomised, Phase 3 Trial.” The Lancet. Oncology 19 (12): 1630–40. https://doi.org/10.1016/S1470-2045(18)30570-9.

Rimawi, Mothaffar F, Rachel Schiff, and C Kent Osborne. 2015. “Targeting HER2 for the Treatment of Breast Cancer.” Annual Review of Medicine 66: 111–28. https://doi.org/10.1146/annurev-med-042513-015127.

Samatar, Ahmed A., and Poulikos I. Poulikakos. 2014. “Targeting RAS-ERK Signalling in Cancer: Promises and Challenges.” Nature Reviews Drug Discovery 13 (12): 928–42. https://doi.org/10.1038/nrd4281.

Sato, Hiroki, Hiromasa Yamamoto, Masakiyo Sakaguchi, Kazuhiko Shien, Shuta Tomida, Tadahiko Shien, Hirokuni Ikeda, et al. 2018. “Combined Inhibition of MEK and PI3K Pathways Overcomes Acquired Resistance to EGFR-TKIs in Non-Small Cell Lung Cancer.” Cancer Science 109 (10): 3183–96. https://doi.org/10.1111/cas.13763.

Satpathy, Shankha, Eric J. Jaehnig, Karsten Krug, Beom Jun Kim, Alexander B. Saltzman, Doug W. Chan, Kimberly R. Holloway, et al. 2020. “Microscaled Proteogenomic Methods for Precision Oncology.” Nature Communications 11 (1). https://doi.org/10.1038/s41467-020-14381-2.

Schmidlin, Thierry, Donna O. Debets, Charlotte A.G.H. van Gelder, Kelly E. Stecker, Stamatia Rontogianni, Bart L. van den Eshof, Kristel Kemper, et al. 2019. “High-Throughput Assessment of Kinome-Wide Activation States.” Cell Systems 9 (4): 366-374.e5. https://doi.org/10.1016/j.cels.2019.08.005.

Schneeweiss, A., S. Chia, T. Hickish, V. Harvey, A. Eniu, R. Hegg, C. Tausch, et al. 2013. “Pertuzumab plus Trastuzumab in Combination with Standard Neoadjuvant Anthracycline-Containing and Anthracycline-Free Chemotherapy Regimens in Patients with HER2-Positive Early Breast Cancer: A Randomized Phase II Cardiac Safety Study (TRYPHAENA).” Annals of Oncology 24 (9): 2278–84. https://doi.org/10.1093/annonc/mdt182.

Spicer, James, Sydonia Rayter, Neville Young, Richard Elliott, Alan Ashworth, and Darrin Smith. 2003. “Regulation of the Wnt Signalling Component PAR1 A by the Peutz-Jeghers Syndrome Kinase LKB1.” Oncogene 22 (30): 4752–56. https://doi.org/10.1038/sj.onc.1206669.

Swain, Sandra M., José Baselga, Sung-Bae Kim, Jungsil Ro, Vladimir Semiglazov, Mario Campone, Eva Ciruelos, et al. 2015. “Pertuzumab, Trastuzumab, and Docetaxel in HER2-Positive Metastatic Breast Cancer.” New England Journal of Medicine 372 (8): 724–34. https://doi.org/10.1056/nejmoa1413513.

Swain, Sandra M., Sung Bae Kim, Javier Cortés, Jungsil Ro, Vladimir Semiglazov, Mario Campone, Eva Ciruelos, et al. 2013. “Pertuzumab, Trastuzumab, and Docetaxel for HER2-Positive Metastatic Breast Cancer (CLEOPATRA Study): Overall Survival Results from a Randomised, Double-Blind, Placebo-Controlled, Phase 3 Study.” The Lancet Oncology 14 (6): 461–71. https://doi.org/10.1016/S1470-2045(13)70130-X.

Tang, Xiaoli, Meiyuan Yang, Zheng Wang, Xiaoqing Wu, and Daorong Wang. 2019. “MicroRNA-23a Promotes Colorectal Cancer Cell Migration and Proliferation by Targeting at MARK1.” Acta Biochimica et Biophysica Sinica 51 (7): 661–68. https://doi.org/10.1093/abbs/gmz047.

Voort, Anna Van Der, Mette S. Van Ramshorst, Erik D. Van Werkhoven, Ingrid A. Mandjes, Inge Kemper, Annelie J. Vulink, Irma M. Oving, et al. 2021. “Three-Year Follow-up of Neoadjuvant Chemotherapy with or without Anthracyclines in the Presence of Dual ERBB2 Blockade in Patients with ERBB2-Positive Breast Cancer: A Secondary Analysis of the TRAIN-2 Randomized, Phase 3 Trial.” JAMA Oncology, 1–7. https://doi.org/10.1001/jamaoncol.2021.1371.

Wagner, Erwin F., and Ángel R. Nebreda. 2009. “Signal Integration by JNK and P38 MAPK Pathways in Cancer Development.” Nature Reviews Cancer 9 (8): 537–49. https://doi.org/10.1038/nrc2694.

Weiner, Timothy M, Edison T Liu, and Rolfj Craven. 1990. “And Invasive Cytogenetic versus DNA Diagnosis In.” Biological Chemistry, no. 2: 1024–25.

